# Data architecture for a large-scale neuroscience collaboration

**DOI:** 10.1101/827873

**Authors:** The International Brain Laboratory, Niccolò Bonacchi, Gaelle Chapuis, Anne Churchland, Kenneth D. Harris, Max Hunter, Cyrille Rossant, Maho Sasaki, Shan Shen, Nicholas A. Steinmetz, Edgar Y. Walker, Olivier Winter, Miles Wells

## Abstract

Effective data management is a major challenge for neuroscience labs, and even greater for collaborative projects. In the International Brain laboratory (IBL), ten experimental labs spanning 7 geographically distributed sites measure neural activity across the brains of mice making perceptual decisions. Here, we report a novel, modular architecture that allows users to contribute, access, and analyze data across this collaboration. Users contribute data using a web-based electronic lab notebook (Alyx), which automatically registers recorded data files and uploads them to a central server. Users access data with a lightweight interface, the Open Neurophysiology Environment (ONE), which searches data from all labs and loads it into MATLAB or Python. To analyze data, we have developed pipelines based on DataJoint, which automatically populate a website displaying a graphical summary of results to date. This architecture provides a new framework to contribute, access and analyze data, surmounting many challenges currently faced by neuroscientists.

## Introduction

Three changes in the field of neuroscience have given rise to new challenges in data management. First, datasets are becoming vastly larger. For instance, advances in neural measurement technology have made it possible to collect neural activity from hundreds of neurons in daily experiments^1,2^. These neural datasets are often accompanied by increasingly large behavioral datasets, often including video data tracking of the animal^3–6^. Second, many neuroscientists are now assembling into teams that include multiple researchers and universities and even spanning several countries^7–10^. This team approach to neuroscience necessitates that data are made immediately accessible to team members at multiple sites, thus demanding a more rigorous data management system than is required in an individual lab. Finally, the field has come to appreciate the need for public data sharing. Data sharing allows a single dataset to generate many insights into brain function, often far beyond those that were the focus of the original research. In order for a dataset to be successfully shared, it must be formatted so that it is readable by new users, must include relevant metadata, and must be stored reliably to ensure access.

Despite the widespread need for improved data management, many neuroscience labs lack the infrastructure needed to improve their current systems. Methods of data storage have not evolved to support the increasingly large datasets being generated. Newly established collaborations struggle to find affordable and efficient ways to pool and visualize data across a large team. Finally, standardized data formats^11,12^ have not been fully adopted, especially compared to data formats in other fields such as genomics and astronomy.

The International Brain Laboratory (IBL) is a collaboration of 21 neuroscience laboratories studying the computations supporting decision-making^7^. The collaboration is generating unprecedented amounts of behavioral and neural data which must be available immediately to all members of the collaboration, and later to the general public. Data management within IBL is made simpler by our initial focus on a single behavioral task implemented with standardized experimental instrumentation^13^. Nevertheless, the scale and diversity of the data produced, and the requirement for immediate access across a geographically-distributed collaboration required us to develop new tools for data management.

An initial goal of the IBL project is to collect a brain-wide activity map: sampling 270 recording sites twice each using Neuropixels probes^1^, along with a single site that is sampled more heavily to assess repeatability across laboratories. This will generate recordings from ~200,000 neurons. In addition to this brain-wide map, individual IBL scientists will perform diverse experiments applying other recording modalities to the same task (e.g. two photon calcium imaging, fiber photometry of neuromodulators), or perform additional manipulations such as optogenetic stimulation or inactivation of specific cell populations. All these data must be put into a standard form and collected into a single central location.

Public release of these data will require careful quality control and manual curation, and will only occur after sufficient time has allowed for this (planned for September 2020). In the meantime however internal IBL scientists require access to these data from multiple international locations. Different scientists require different data items, to perform tasks such as developing quality control metrics and analyzing behavioral learning curves. Data sharing solutions that require researchers to download the entire dataset to access specialized data items would therefore be impractical, adding unnecessary obstacles to the already difficult task of analyzing and exploring complex datasets. Furthermore, scientists need to be able to search for experiments and data items matching specified criteria. We have now established our core data pipeline and sharing architecture, which solves these problems, and will also form the basis of our external sharing system when the data are publicly released. As of January 2020, this system stores information on 16,000 behavioral sessions (currently 400,000 files).

Here, we describe the methods and open-source software IBL has developed to contribute, access and analyze data. This data architecture is composed of three modules (Figure 1), which are described in detail below. The first, “Alyx,” allows users to contribute data via a web-based electronic lab notebook system. Alyx stores metadata concerning experimental subjects, experiments, and data files, and automatically transfers recorded data to a single central location. The second, “Open Neurophysiology Environment” is a simple, flexible, and extensible protocol for accessing data files. Finally, a data pipeline based on Datajoint^14^, automatically analyzes incoming data, and powers a web viewer for monitoring behavioral performance of mice, and viewing activity of individual neurons. These modules are all available as open source, and can be adopted individually or together by other projects, as their needs require.

**Figure 1.**
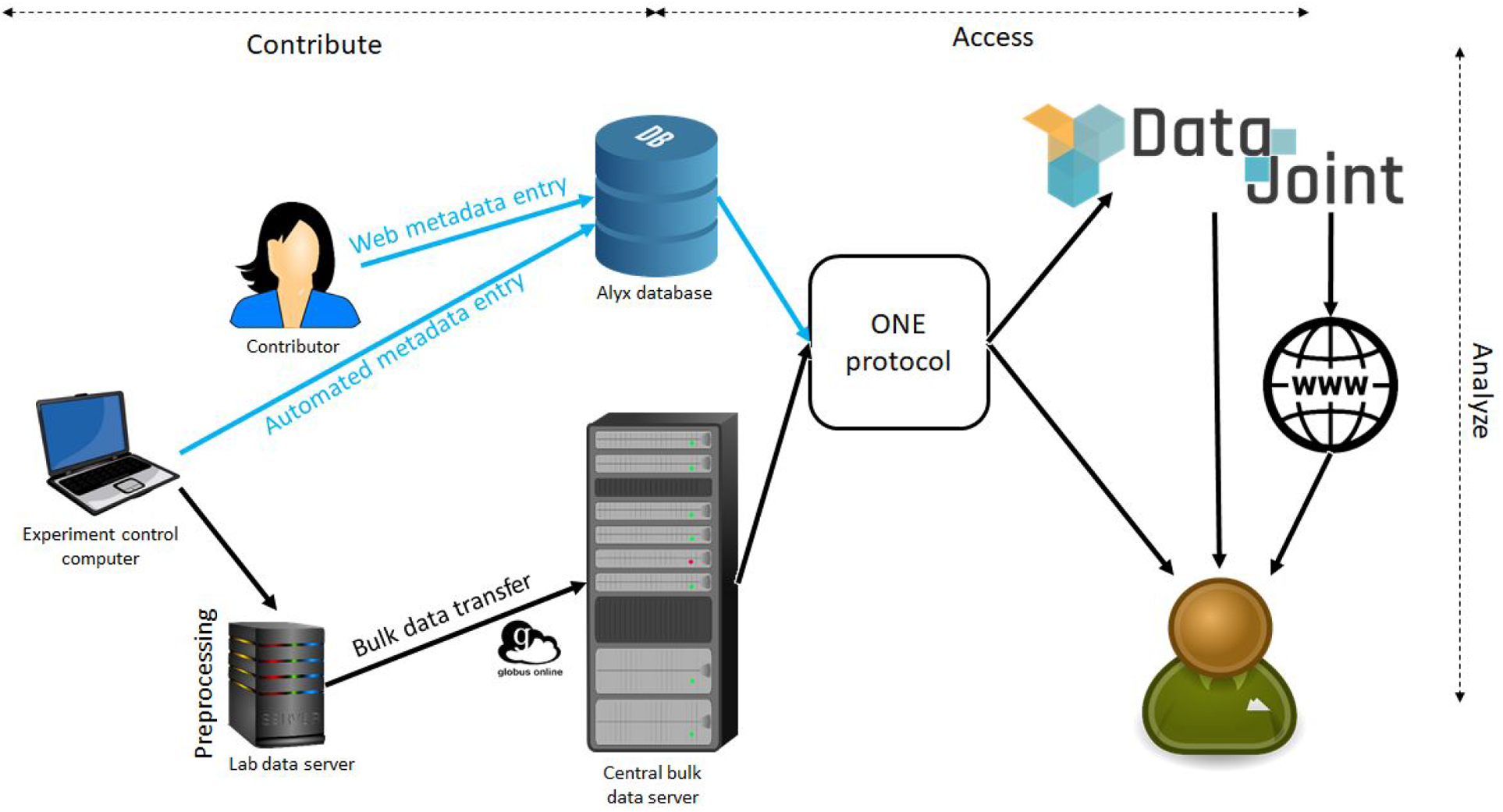
IBL data architecture. Data and metadata contributed from each lab are integrated into a relational database storing metadata, and a bulk file server storing binary files, via read-only connections. Users can access these data either directly through the Open Neurophysiology Environment (ONE) protocol, or via DataJoint, which also allows automatic pipelined analysis, and presents basic analyses on a website.

## Contributing data: an electronic lab notebook for metadata

A key challenge is to ensure that metadata (Table 1) are comprehensive and accurate. While metadata about the experiments themselves can be collected by the recording hardware, metadata about the experimental subjects must be entered by lab members. Ensuring this happens reliably is primarily a problem of “social engineering” rather than software, and we have solved it by creating a user-friendly web-based client that connects from all IBL labs to our central database, termed “Alyx” (Fig. 1). Alyx is accessible via a web interface that allows data entry for example real-time note taking during surgery or experiments. It is also programmatically accessible via an API. This allows the experimental control software to communicate directly with the database, so metadata such as experiment time, the amount of water delivered, as well as the locations and types of all recorded data files, can be automatically registered.

**Table 1:**
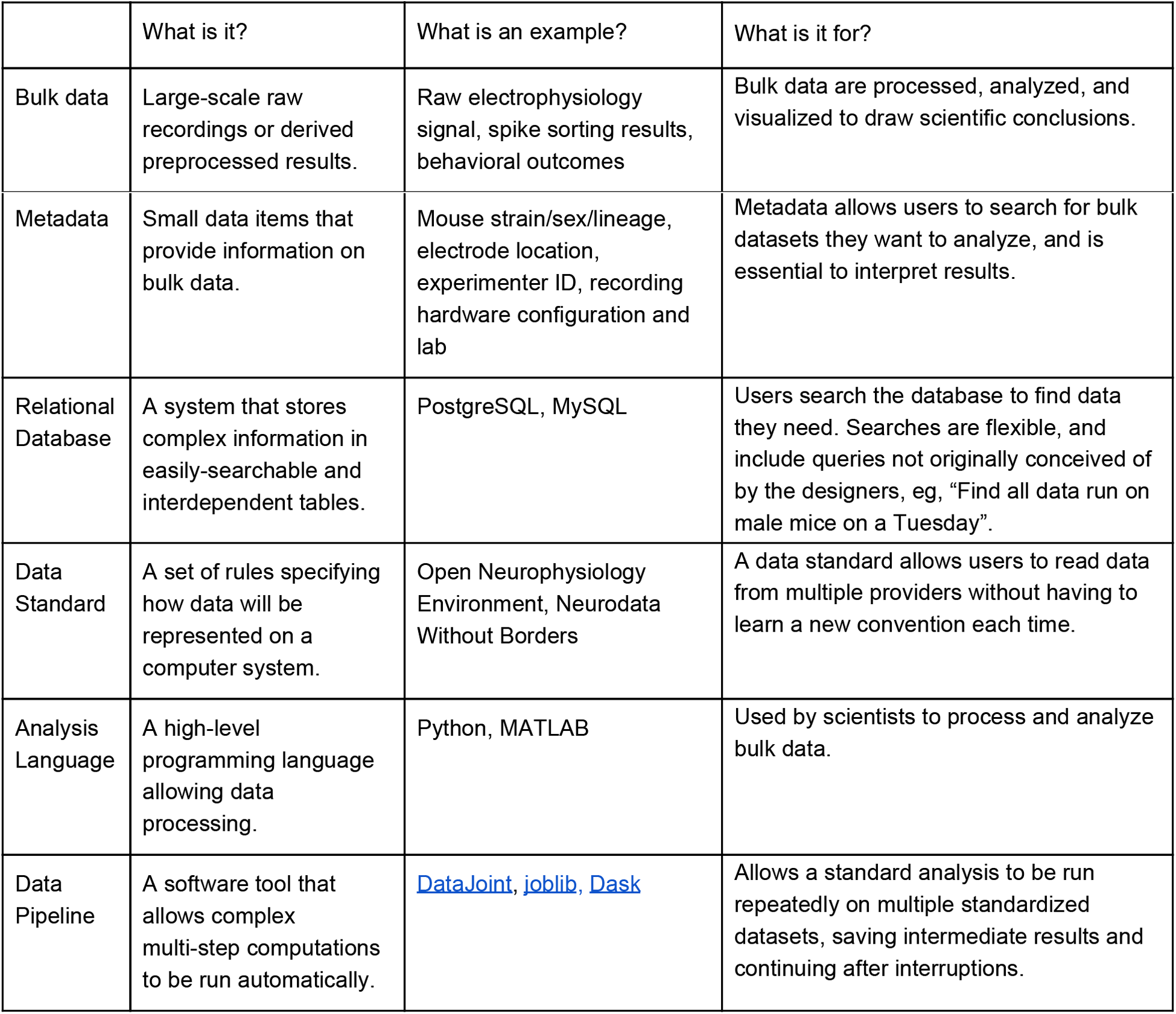
Terminology

The Alyx system functions as an electronic colony management / electronic lab notebook system storing details on all labs’ mice, such as age and surgical history. These are critical because mouse behaviour is affected by a large number of factors^15^. Alyx allows metadata to be entered at the time of collection, for example recording each subject’s weight before every experiment, which ensures data are entered more reliably than would occur by transcribing from paper lab notebooks. The system also performs other functions, such as generating email notifications telling the experimenter how much supplementary water is needed after training, and recording delivery of these supplements. Other types of metadata (e.g. genotype results), are entered as soon as they are generated, which ensures their accuracy, in contrast to later entry of paper-based notes. Preprocessing software (eg. spike sorting, compression) also communicates with the database to register the files it has produced. This registration of files enables automatic transfer of data to the central server.

Alyx can function as a system for colony management and registration of experiments and data, independently of the remainder of the IBL data architecture. Indeed, it was first developed by one of the IBL member labs for this purpose prior to IBL’s founding, and is now used by several other labs to perform colony management for non-IBL mice. It can be installed on a local computer or on a remote cloud server (eg. AWS) accessed by web interface. A user guide to Alyx is available as Appendix 5 of this paper.

## Contributing data: preprocessing and transfer of bulk data

The experimental data are generated via neural and behavioral measurements during a dynamic decision-making task for mice^7^. Data are collected daily at multiple experimental sites, during both initial training and neurophysiological recordings in expert subjects. In addition to neural recordings, these data include the output of rotational sensors on the wheel that subjects use to report choices, video recordings of the face and body from 3 high-speed cameras, a microphone recording of sound inside the apparatus, and measurements from additional sensors that track ambient environmental parameters (e.g. temperature).

Preprocessing and compression run on a local data server in each experimental lab. This server runs on a widely used Linux distribution and is assembled using consumer grade components: gaming GPU, 64 Gb of RAM, solid state system drive plus extra hard drives for temporary storage. Prior to upload, to save space and bandwidth on the bulk dataserver. Video is compressed using ffmpeg h264 codec (compression level crf 29), and tracking of body parts is performed with DeepLabCut^4^ Raw electrophysiology is compressed by a factor of ~3 using a custom lossless compression algorithm we have developed specifically for this purpose (https://github.com/int-brain-lab/mtscomp; Appendix 3). Spike sorting is performed using kilosort2 (https://github.com/MouseLand/Kilosort2). A versioning system ensures that the software version used to generate these processed files is recorded, so future improvements in preprocessing will be handled gracefully. The data produced by these experiments are stored primarily as .npy files - a simple format consisting of flat binary data plus a small header. These files can be natively accessed by Python, and can be accessed by MATLAB using a simple interface (https://github.com/kwikteam/npy-matlab). Files are named using a convention based on ONE dataset types (Appendix 1).

Data are automatically uploaded to the bulk data server (Table 1, Fig. 1, middle) via Globus Online, a system that allows transfer of large data sets, allowing for features such as automatic restart and parallelization over multiple Transmission Control Protocol (TCP) connections. This transfer is initiated at 2am local time by the central Alyx database server. The database keeps a record for each dataset including the version and the file hash, plus an additional linked record for each physical copy of a dataset. Any datasets whose presence is registered as residing on the lab server, but not the central server, are uploaded and their presence on the central server is registered when the upload completes.

The IBL experiment control software, including software for automatically training mice, as well as all hardware designs are open source and available for scientists who wish to study this behavior task (https://www.internationalbrainlab.com/resources). The software for data compression and transfer are open source (https://www.internationalbrainlab.com/resources), with remote data transfer also requiring an account with Globus Online.

## Accessing the data

Data on the central server is standardized and shared within the collaboration using the Open Neurophysiology Environment (ONE), a simple protocol developed for this purpose (Fig. 1; https://ibllib.readthedocs.io/en/latest/04_reference.html#open-neurophysiology-environment).

ONE is a development from the Neurodata Without Borders (NWB) standard^11,12^, that adds additional features required for use in a distributed collaboration such as IBL. A Jupyter hub allowing public access to the behavioral data of the companion paper^13^ via ONE is available at https://one.internationalbrainlab.org.

ONE comprises a set of conventions for naming the datasets associated with an experiment, and a lightweight interface allowing users to search for experiments of interest, and load required data from these experiments directly into in MATLAB or Python (Appendix 1). The user thus need not worry about underlying file formats or network connections, and data are cached on their local machine to avoid repeated downloads.

The key to ONE’s standardization is the “dataset type”. When a user loads data from one of these types, they are guaranteed to be returned predictable information, organized in a predictable way, and measured in a predictable unit. For example, if a user requests the dataset `spikes.times` for a particular experiment, they are returned the times of all extracellularly recorded spikes, measured in seconds relative to experiment start, formatted as a 1-dimensional column vector. The current list of ONE-standard dataset types is an extension of the NWB data schema, and is shown in Appendix 2. Datasets need not be two-dimensional numeric arrays – they can be arrays of any dimensionality, lists of strings, movies, or arrays of structures. The user is guaranteed when loading a dataset of a given type that they will be returned data organized the same way, regardless of which experiment they load from.

Dataset types are named in a specific way, formalizing relations between data. Each dataset type has a two-part name: the “object” (before the period) and the “attribute” (after the period). Multiple dataset types with the same object (e.g. `spikes. times` and `spikes. clusters`), always have the same number of rows, describing multiple attributes of the same object: in this case, the times and cluster assignments of each spike. If the attribute of one dataset matches the object of another, this represents a cross-reference. For example, `spikes.ciusters` contains an integer cluster assignment for each spike, while `ciusters.brainLocation` contains a structure array with information on the physical location of each of these cells. The values in `spikes.clusters` can be used as indices to the rows of `clusters.brainLocation`.

ONE was designed to be an extensible data standard (Table 1) that can be used beyond IBL, allowing other data providers to extend it as required while still maintaining compatibility. Data that are expected to be common across projects are encoded in standard dataset types (such as `spikes.times`), and all providers of data containing these types should return arrays of the specified dimension and measurement units. However, providers can also add their own project-specific dataset types, by defining their own “namespace”. Names beginning with an underscore are guaranteed never to be standard. For example the dataset type `_ibl_trials.stimulusContrastLeft` contains information specific to the IBL project, and no other projects would be expected to use it.

ONE can be used as a stand-alone module, independently of the full IBL data architecture. To enable users to implement ONE without the rest of the architecture, we have provided an implementation termed ONE light (https://github.com/int-brain-lab/ibllib/tree/onelight/). To share data with ONE light, a data provider gathers their data files into experiment directories, with files named according to the standard dataset types. They then run a script that automatically uploads the files to figshare. This system has been used to share large-scale electrophysiology recordings collected in a different task^16^ (https://figshare.com/articles/steinmetz/9974357). A user can access this data using the same standard library functions as for the main IBL dataset. ONE light can also be used to access data stored on a user’s local machine, and we have provided access to IBL’s behavioral data using this system (https://figshare.com/articles/A_standardized_and_reproducible_method_to_measure_decisionmaking_in_mice_Data/11636748).

Once curation is complete, data will also be made available via Neurodata Without Borders (NWB)^11,12^. NWB was created by neurophysiologists and software developers to be a unified data standard (Table 1) suitable for diverse neurophysiological and behavioral data. NWB can store multimodal experimental data in a single file and thus is well suited for long-term distribution and storage. ONE has additional features that make it suitable for a growing data set with frequent contributions and diverse access needs (for example, the ability to load a single data item such as behavior performance from all experiments without downloading all full data files). ONE’s standardized data access functions will facilitate production of NWB files, and will also allow flexibility as neurodata standards evolve in the future. Release of data in NWB format has been successfully piloted by individual laboratories within the collaboration^17,18^.

## Analyzing data

The final part of the data architecture is the system for analyzing data collected by the project. This component of the architecture is built on DataJoint^14^: a workflow management system that integrates a relational database (Table 1) with automatic processing of analyses. While DataJoint and Alyx are both based on relational databases, the design philosophy and intended uses of these systems is different, and they are hosted on different servers. Unlike Alyx, whose table structure is not designed to change, DataJoint is a dynamic system that allows individual users to create and delete new tables containing results of ad-hoc analyses. It thereby allows users to rapidly explore new analysis methods, and easily scale them up to run on the entire IBL data set. Furthermore, because all computation results are stored in the database, the system allows users to quickly build secondary analyses making use of the results of previous computations. The analyses in the companion paper^13^ were made using this system, and a Jupyter hub allowing public access to IBL data via DataJoint, together with sample code for some of these figures, is available at https://jupyterhub.internationalbrainlab.org.

In addition to allowing dynamic access to data, we have used the DataJoint system to build a web system for monitoring the behavioral progress of all mice. As new data are recorded, they are automatically ingested into the DataJoint system, and analyses summarizing the behavioral performance of all mice are automatically updated. Figures summarizing mouse progress are produced, and made available via a web site (Fig. 2). This web system is used by team members who are training mice to monitor progress, or to compare the performance of many animals, and also allows newcomers to the group to quickly get a sense of the behavioral data without needing to write their own analysis routines. A curated subset of these behavioral data are available on a public website (http://data.internationalbrainlab.org).

**Figure 2:**
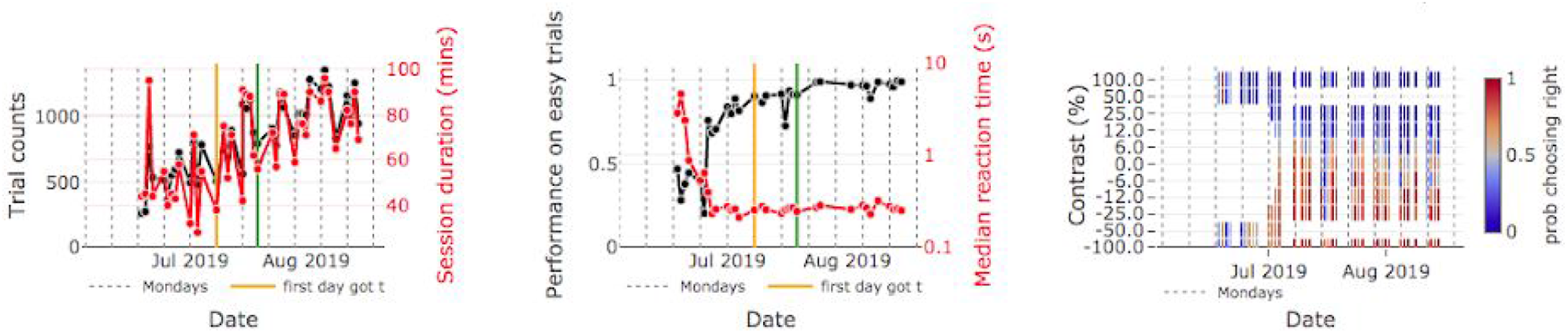
An analysis pipeline (created through DataJoint) automatically generates plots of behavioral data each time a subject is run. Plots generated by this pipeline are instantly available to all team members via a web browser. Data from a single mouse are shown. Left: daily trial counts and session duration; middle: Proportion correct and reaction time on high contrast (easy) trials; Right: Accuracy for all contrast levels over a training period. Note that early training days (left) include only high contrast trials while later training days (right) include a mix of contrasts.

The data analyses displayed on the website are performed using a growing toolbox of standard and emerging analyses for analysis of electrophysiological and behavioral data. The toolbox, termed Brainbox, is written in Python and can operate independently of DataJoint. These tools are publicly available at https://github.com/int-brain-lab/ibllib/tree/master/brainbox, and can be used by neuroscientists independently from the rest of the data architecture.

## Outlook

Data management, standardization and sharing has proven challenging within neuroscience. Datasets have grown in size, complexity and diversity in recent years, but few neuroscientists have prior experience or formal training in data management. Although the IBL is still at an early stage, our experience thus far has shown it is possible to develop a reliable, easy-to-use data architecture, that generates and organizes data suitable for sharing within a large collaboration, and beyond. The IBL, as a large collaboration, has benefitted from outstanding technical staff who work full time on data management, but most individual labs do not have the resources to do this. As a result, data management problems slow down projects, data sharing has not been universally adopted, and many shared datasets are lacking metadata. In this regard, neuroscience is substantially behind other scientific domains, such as genomics, astronomy, or particle physics.

We end with some thoughts on how the lessons we have learned in IBL could inform data management practices in individual labs. One possibility is for individual labs to nominate individuals to focus on data management. However, the time commitment required should not be underestimated: organizing, documenting, sharing, and supporting the data collected in even a medium-sized lab that uses modern neurophysiological methods is essentially a full-time job. An alternative possibility is for labs to rely on third-party companies that offer data management solutions (such as DataJoint Neuro, who have partnered with IBL as well as several individual labs outside the collaboration). In addition to the cost, however, this solution might not appeal to users hesitant to fully rely on a third party, fearing that this will limit flexibility and the ability to nimbly update workflows.

We hope that the open-source tools developed by IBL will help contribute to a solution allowing individual labs to curate and share data, without requiring dedicated full-time staff or external contractors. Our Alyx colony management system provides a solution to the problem of contributing and storing metadata. The ONE Light protocol allows data producers to “upload and forget” datasets at the time of publication, while allowing users a standardized and searchable protocol to access data from multiple producers. The Brainbox library contributes a growing set of analysis tools, and our DataJoint pipeline provides a template data providers can adapt to provide web-based data visualization. We are keen to support the use of these tools by non-IBL labs, so we can continue to to make them more useful and easy to apply by the general community. Our hope is that the IBL’s data management plan can help pave the way forward to a new era of large scale data in neuroscience in which data from all labs is shared on a routine basis.

## Supporting information

Appendix 5: Alyx User Guide

## Author contributions

Niccolò Bonacchi [Conceptualization] (supporting). [Data curation] Data, metadata, and pipeline (equal). [Funding acquisition] (supporting). [Project administration] Meeting coordination (supporting), attendance (equal). [Resources] Computing and data storage (equal). [Software, Validation] Pipeline, core, quality control (equal), database and analysis libraries (supporting). [Visualization] Posters (lead), presentations (equal). [Writing – original draft] (supporting). [Writing – review & editing] (supporting).

Gaelle Chapuis [Data Curation] Helped with writing user guides detailing how to enter metadata in Alyx (supporting). [Software, Validation] Acted as naive user tester for ONE and DJ (supporting). [Project administration] Gathered and reported users’ requirements (supporting).

Anne Churchland Contributed to project administration, funding acquisition, writing and revising.

Kenneth D. Harris. Contributed to design of overall data architecture, and to Alyx and ONE systems. Contributed to project administration, funding acquisition, writing and revising.

Max Hunter. Contributed to the design and development of Alyx.

Cyrille Rossant. Contributed to design and implementation of overall data architecture, and to Alyx and ONE systems.

Maho Sasaki Contributed to the design and implementation of the IBL Data Portal website.

Shan Shen Contributed to the design and implementation of the DataJoint pipeline and the IBL JupyterHub

Nicholas A. Steinmetz contributed to the design, testing, and development of Alyx, of dataset types, and of software tools for working with them.

Edgar Y. Walker Contributed to the design and implementation of the DataJoint pipeline and the IBL Data Portal.

Olivier Winter [Software] Implemented the backend data infrastructure. [Validation/Methodology] Designed full loop integration tests to allow maintenance of the codebase. [Data Curation] Fixed and updated erroneous datasets.

Miles Wells. Contributed to the design, testing and implementation of Alyx, its dataset types and of software tools that work with them.

## Appendix 1: Open Neurophysiology Environment details

The Open Neurophysiology Environment (ONE) user interface consists of four simple API functions that allow data consumers to search for experiments of interest and load data from them. This interface allows multiple backend instantiations, so data consumers can run the same exact code to process data from multiple data producers, even if these different producers store data in different locations. The main IBL data is provided via an ONE implementation that requires a backend SQL database, but we have also provided an “ONE light” implementation that allows data producers to share data simply by uploading files to a web server, or to figshare (a site offering free hosting for scientific data). The same implementation also supports data stored in a local file system, as long as the hierarchical organization into subfolders and file names matches the same ONE convention. These files can be uploaded in a variety of standard formats (npy, csv, json, hdf5, mpeg, etc.), organized with one directory per experiment containing appropriately-named data files.

ONE clients have been implemented in Python and MATLAB, and are documented at https://ibllib.readthedocs.io/en/develop/index.html. Below, we describe how to use ONE in Python.

### Importing ONE

Users can access data from different projects by loading and configuring appropriate implementations of the ONE library. For example, to access IBL’s main data in Python they would type

~~~
from oneibl.one import
ONE one = ONE()
~~~

whereas to access data from Steinmetz et al.^16^ via the ONE light figshare interface, they would type

~~~
from oneibl.onelight import ONE
one = ONE()
one.set_figshare_url(“https://figshare.com/articles/steinmetz/9974357”)
~~~

In both cases, the user is returned an object named one, that allows the same commands to be run whichever ONE implementation is running behind the scenes.

### Dataset types

The key to ONE’s standardization is the “dataset type”. When a user loads data from one of these types, they are guaranteed to be returned predictable information, organized in a predictable way, and measured in a predictable unit. For example, if a user requests the dataset `spikes.times` for a particular experiment, they are guaranteed to be returned the times of all extracellularly recorded spikes, measured in seconds relative to experiment start, and returned as a 1-dimensional column vector. The current list of ONE-standard dataset types was derived from the NWB data schema, and is shown in Appendix 2.

Datasets need not be two-dimensional numeric arrays – they can be arrays of any dimensionality, lists of strings, movies, or arrays of structures. In fact, a dataset can by anything with the concept of a “row” – i.e. any data structure that can be addressed by an integer subscript, for example a leading dimension of a structure array, or frame number in a movie.

Dataset types are named in a specific way, which formalizes relations between data. Each dataset type has a two-part name, with the part before the period called the “object” and the part after called the “attribute”. When there are multiple dataset types with the same object (e.g. `spikes.times` and `spikes.clusters`), it is guaranteed that they will have the same number of rows, describing multiple attributes of the same object: in this case, the times and cluster assignments of each spike. If the attribute of one dataset matches the object of another, this represents a cross-reference. For example, `spikes.clusters` contains an integer cluster assignment for each spike, while `clusters.brainLocation` contains a structure array containing information on the physical location of each of these cells. The values in `spikes.clusters` can be used as indices to the rows of `clusters.brainLocation`.

Dataset types have specified measurement units. For example, locations in the brain are given in Allen CCF coordinates^19^, measured in mm. The units of measurement for all dataset types are specified in the documentation.

Some attributes have predefined meanings. For example it is guaranteed that any dataset type whose attribute is of the form `times` or `*_times` will represent the times of events, measured in seconds relative to experiment start. Datasets whose attribute is `intervals` or `*_intervals` are guaranteed to be two-column arrays giving start and end times of particular events in seconds relative to experiment start. Datasets whose attribute is `timestamps` contain timestamps for each sample of not-necessarily evenly sampled timeseries data, whose unit is again seconds relative to experiment start.

Not all data can be standardized. Data that are common across projects will be encoded in standard dataset types, and providers of data containing these types should return arrays of the specified dimension and measurement units. However, providers can add their own project-specific dataset types, by defining their own “namespace”. Names beginning with an underscore are guaranteed never to be standard. For example the dataset type `_ibl_trials.stimulusContrastLeft` contains information specific to the IBL project, and no other projects would be expected to use it. A list of standard dataset types will be maintained centrally, and will start small but increase over time as the community converges on good ways to standardize more information. The current list of standard and IBL-specific dataset types is in appendix 2.

### Searching for data

Each experiment is characterized by a unique string known as the experiment ID (eID). The format of this string is determined by the implementation: ONE light uses directory pathnames, while the main IBL implementation uses hexadecimal UUIDs. Within an implementation, a specific eID is always guaranteed to refer to the same experiment, allowing data consumers to hard-code eID strings into their analysis software.

To search for experiments they want to analyze, a user runs the function `one.search` that returns a list of eIDs for all experiments matching the requested criteria. ONE currently allows searching for experiments by user (i.e. experimenter); by experimental subject; by date; and by the dataset types associated with the experiment. For example, to search for all experiments on subject ‘IBL_123’ performed in the month of August 2019, for which spike-sorted extracellular electrophysiology and video eye tracking are available, one would type

~~~
eID list = one.search(subject=‘IBL 123’, date range=[‘2019-08-02’, ‘2019-08-31’], dataset types=[‘spikes.times’, ‘spikes.clusters’,eye.xyPos’])
~~~

This command returns a list of eIDs for all experiments matching the specified criteria. Optionally, the search command also returns full metadata on each matching experiment in a structure list.

### Loading data

Once a user knows the eID of an experiment they want to analyze, they can load data from this experiment in one of two ways. The first way is to load an individual dataset. For example, to load spike times, the user would type:

~~~
st = one.load dataset(eID, ‘spikes.times’)
~~~

The second way is to load all datasets belonging to an object. For example the command

~~~
spikes = one.load object(eID, ‘spikes’)
~~~

will return a dictionary with one entry for each dataset associated with that object (`times, clusters, depths, amps`, etc.).

Importantly, the user need not be concerned with the physical location or format of the raw data files - they just type these commands to load them into their analysis software. In practice, the data are cached on the user’s local machine, so it need only be downloaded once for many uses; however details of the caching, as well as the underlying file formats are hidden from the user.

### Listing available data

The fourth and final ONE function `one.list(eID)` simply lists the dataset types available for a given experiment.

## Appendix 2: ONE dataset types

The table below lists the dataset types currently used in the IBL’s implementation of the ONE standard. The standard namespace contains data we believe can be standardized with other projects; these are largely adopted from the NWB data model. The _ibl_ namespace contains data likely to be specific to our task or recording hardware. A live list of dataset types is maintained at https://www.internationalbrainlab.com/resources. The dataset types as of the time of writing are listed below.

We have provided a “ONE Light” interface, that allows any scientists to share data via ONE protocols, without needing to run a backend database. To do so, they create a data directory for each experiment, containing one data file for each of the dataset types they wish to provide. The files can be in multiple formats, but the standard is .npy files: flat binary files encoding numerical arrays, with a small header indicating the size and datatype. These files can be loaded and saved natively from numpy, and from MATLAB via <u>this toolbox</u>. For example, to store spike times, the experiment directory would contain a file “spikes.times.npy”. Once a data sharer has created directories are creating containing these data files on their local machine, they run a command using <u>this library</u> to automatically upload them to figshare or to a web server. Data users can then search and download data using the standard ONE interface, though currently with simpler search capabilities than with the backend database.

| ONE Dataset Type | Dimension | Description |
| --- | --- | --- |
| spikes.times | [nspi] | Times of spikes (seconds, relative to experiment onset). Note this includes spikes from all probes, merged together. |
| spikes.clusters | [nspi] | Cluster assignments for each spike (integers counting from 0). Cluster assignment reflects the result of manual curation. |
| spikes.depths | [nspi] | Depth along probe of each spike ($\mu\text{m}$; computed from waveform center of mass). 0 means deepest site, positive means above this. |
| spikes.amps | [nspi] | Peak amplitude of each spike ($\mu\text{V}$). |
| spikes.templates | [nspi] | Template ID for each spike as originally assigned by spike sorting software, prior to manual curation. |
| spikes.samples | [nspi] | Time of spikes, measured in units of samples in their own electrophysiology binary file. |
| templates.waveforms | [ntemp, nsw, nchSub] | Waveform of each template spike (stored as a sparse array, only for a subset of channels with large waveforms). |
| templates.waveformsChannels | [ntemp, nchSub] | Channels of the raw recording on which the template waveforms are defined. |
| clusters.metrics | [nc, nmetrics] | Quality control metrics for each cluster. |
| clusters.mlapdv | [nc, 3] | Estimated coordinates of the cell relative to bregma (mm) in Allen Common Coordinate Framework (CCF)<sup>15</sup>. See Appendix 4. |
| clusters.brainAcronyms | nc | Estimated cluster brain location acronym using Allen CCF |
|  |  | notation. See Appendix 4. |
| clusters.waveforms | [nc, nsw, nchSub] | Mean unfiltered waveform of spikes in this cluster (NB: Neuropixels 1.0 data will have been hardware filtered). |
| clusters.waveformsChannels | [nc, nchSub] | Identities of the channels that are represented in clusters.meanWaveforms for each cluster sorted by amplitude. |
| clusters.depths | [nc] | Depth of mean cluster waveform on probe ($\mu\text{m}$). 0 means deepest site, positive means above this. |
| clusters._phy_annotation | [nc] | Manual curation of spike cluster quality. 0 = noise, 1 = MUA, 2 = Good, 3 = Unsorted, 4-100 = manual quality score. |
| clusters._phy_ids | [nc] | Original cluster index assigned in Phy (counting from 0). |
| clusters.peakToThrough | [nc] | Trough to peak time of mean cluster waveform (ms). |
| clusters.amps | [nc] | Mean amplitude of each cluster ($\mu\text{V}$). |
| clusters.channels | [nc, nch] | Channel which has the largest amplitude for this cluster. NB this counts all channels from all probes; to find this cluster's brain location index like so:<br>channels.brainLocation[clusters.peakChannel[i],:] |
| clusters.probes | [nc, np] | Which probe this cluster came from (counting from zero). |
| probes.trajectory | [np, 7] | Trajectory coordinates of probe. See Appendix 4. |
| probes.description | [np] | Text description of probe: label (folder name), Model (3A, 3B1, 3B2), Serial Number, Original file name. |
| channels.probes | [nch] | Probe assignments for each channel (integers counting from 0). Can be used as direct indexing for the probes.* attributes. |
| channels.rawInd | [nch] | Array of indices in the raw recording file (of its home probe) that each channel corresponds to (counting from zero). |
| channels._phy_ids | [nch] | Array of indices in Phy that each channel corresponds to (counting from zero). |
| channels.brainAcronyms | [nch, 4] | Channel brain location acronym using Allen CCF notation. |
| channels.mlapdv | [nch, 3] | Channel location relative to bregma (mm) in Allen CCF. See Appendix 4. |
| channels.localCoordinates | [nch, 2] | Location of each channel relative to probe coordinate system ($\mu\text{m}$): x (first) dimension is on the width of the shank; (y) is the depth where 0 is the deepest site, and positive above this. |
| channels.shank | [nch] | Array giving shank number (or other channel group) of each channel, used for spike sorting. |
| eye.timestamps | [nEyeSamples, 2] | Timestamps for pupil tracking timeseries: 2 column array giving sample number and time in seconds. |
| eye.raw | [nEyeSamples, nX, nY] | Raw movie data for pupil tracking. |
| eye.area | [nEyeSamples] | Area of pupil (pixels^2). |
| eye.xyPos | [nEyeSamples, 2] | Matrix with 2 columns giving x and y position of pupil (in pixels). |
| eye.blink | [nEyeSamples] | Boolean array saying whether eye was blinking in each frame. |
| licks.times | [nLicks] | Times of licks in seconds. |
| spontaneous.intervals | [nSpontInt, 2] | Times when no other protocol was going on for at least 30 seconds. |
| _ibl_wheel.position | [nWheelSamples] | Absolute linear displacement of wheel (cm). |
| _ibl_wheel.timestamps | [nWheelSamples, 2] | Times of position in seconds relative to session start, continuous (evenly spaced). |
| _ibl_wheel.velocity | [nWheelSamples] | Tangential velocity of the wheel (cm/s) where positive = CW. |
| _ibl_wheelMoves.intervals | [nWheelMoves, 2] | 2 column array of onset and offset times of detected wheel movements in seconds relative to session start. |
| _ibl_wheelMoves.type | [nWheelMoves] | String array containing classified type of movement ('CW', 'CCW', 'Flinch', 'Other'). |
| _ibl_trials.intervals | [nTrials, 2] | Start (i.e. beginning of quiescent period) and end (i.e. end of iti) times of each trial in seconds relative to session start. |
| _ibl_trials.included | [nTrials] | Boolean array of which trials to include in analysis, chosen at experimenter discretion, e.g. by excluding the block of incorrect trials at the end of the session when the mouse has stopped. |
| _ibl_trials.repNum | [nTrials] | The trial repetition number, i.e. how many trials have been repeated on this side (counting from 1). |
| _ibl_trials.goCue_times | [nTrials] | Time of go cue tone in seconds relative to session start. |
| _ibl_trials.goCueTrigger_times | [nTrials] | Time of go cue trigger command was sent in seconds relative to session start. |
| _ibl_trials.response_times | [nTrials] | Time in seconds relative to session start when a response was recorded (end of the closed loop state in bpod). |
| _ibl_trials.choice | [nTrials] | The response ID: -1 (turn CCW), +1 (turn CW), or 0 (nogo) |
| _ibl_trials.stimOn_times | [nTrials] | Times of stimulus onset in seconds relative to session start. |
| _ibl_trials.stimOnTrigger_times | [nTrials] | Times of stimulus onset trigger command in seconds relative to session start. |
| _ibl_trials.contrastLeft | [nTrials] | Contrast of left-side stimulus (0-1, nan if stimulus is |
|  |  | on the other side). |
| _ibl_trials.contrastRight | [nTrials] | Contrast of right-side stimulus (0-1, nan if stimulus is on the other side). |
| _ibl_trials.feedback_times | [nTrials] | Time of feedback delivery (reward or noise) in seconds relative to session start. |
| _ibl_trials.feedbackType | [nTrials] | Whether feedback is positive or negative (-1 for negative, 1 for positive, 0 for no feedback). |
| _ibl_trials.rewardVolume | [nTrials] | Volume of reward given each trial (μl). |
| _ibl_trials.itiDuration | [nTrials] | Inter-trial interval duration for each trial (seconds). |
| _ibl_trials.deadTime | [nTrials] | Time between state machine trial end and restart for every trial. |
| _ibl_trials.probabilityLeft | [nTrials] | Probability (0-1) that the stimulus will be on the left-hand side. |
| _ibl_passiveTrials.included | [nPassiveTrials] | Boolean suggesting which passive trials to include in analysis, chosen at experimenter discretion. |
| _ibl_passiveTrials.stimOn_times | [nPassiveTrials] | Times of stimulus onset in seconds relative to session start. |
| _ibl_passiveTrials.contrastLeft | [nPassiveTrials] | Contrast of left-side stimulus (0-1, nan if stimulus is on the other side). |
| _ibl_passiveTrials.contrastRight | [nPassiveTrials] | Contrast of right-side stimulus (0-1, nan if stimulus is on the other side). |
| _ibl_passiveValveClicks.times | [nPassiveTrials] | Times of valve opening during passive trial presentation in seconds relative to session start. |
| _ibl_passiveBeeps.times | [nPassiveTrials] | Times of the beep, equivalent to the go cue during task, in seconds relative to session start. |
| _ibl_passiveWhiteNoise.times | [nPassiveTrials] | Times of white noise bursts, equivalent to the negative feedback sound during the task, in seconds relative to session start. |
| _ibl_passiveNoise.intervals | [nPassiveTrials,2] | Passive noise trial start and end (i.e. end of iti) times in seconds relative to session start. |
| _ibl_sparseNoise.xy | [nSparseNoise,2] | 2 column array giving x and y coordinates on screen of sparse noise stimulus squares. |
| _ibl_sparseNoise.times | [nSparseNoise] | Times of sparse noise stimulus onset in seconds relative to session start. |
| _ibl_extraRewards.times | [nExtraRewards] | Times of extra rewards in seconds relative to session start. |
| camera.dlc | [nframes, npoints x 3] | Coordinates of DeepLabCut (DLC) points (x position, y position, likelihood). Total points = 19 (fingers-8, nose-2, spout-2, tongue-2, eye-4). |
| camera.times | nframes | Time of each frame acquisition (leftCamera only for behavior, and right and bodyCameras for ephys rigs). |

## Appendix 3: Lossless compression algorithm

Our aim in developing the compression algorithm was to reduce data size, while maintaining not only the full signal available in the original data, but also maintaining the ease of random-access available with flat binary files. To do this, we took advantage of the temporal correlations in electrophysiological recordings, which show an approximate 1/f power spectrum.

The input to the algorithm is represented as a flat binary multiplexed file of 2-byte integers. Data are compressed independently in consecutive chunks of one second, which allows random access to any part of the recording without decompressing the whole signal. To compress a chunk, we first compute discrete time differences independently for each channel, which approximately whitens the signal. We then compress the result using the zlib lossless compression algorithm. The initial values for each chunk and compressed difference signals are then appended to a compressed binary file on a chunk-by-chunk basis, and a companion JSON file is saved storing the byte offset of every chunk. A decompression algorithm reads the JSON and binary files allowing random “slices” of the data to be retrieved on the fly without decompressing the whole file. The compression code is unit-tested with 100% coverage. In our benchmarking we could achieve a ~3x compression ratio of our data. For our ~400 channel recordings, compression is ~4x faster and decompression is ~3x faster than real time on a Intel i9 10-core computer. To the best of our knowledge, this simple compression algorithm has not been previously described, and could be used in other applications that rely on multichannel time series of approximate 1/f spectrum, within and beyond neuroscience.

## Appendix 4: Coordinate system within Allen CCF

Bregma is defined as Voxel ML-566, AP-540, DV-33 within the 10μm volume of the Allen CCF mouse Atlas^19^.

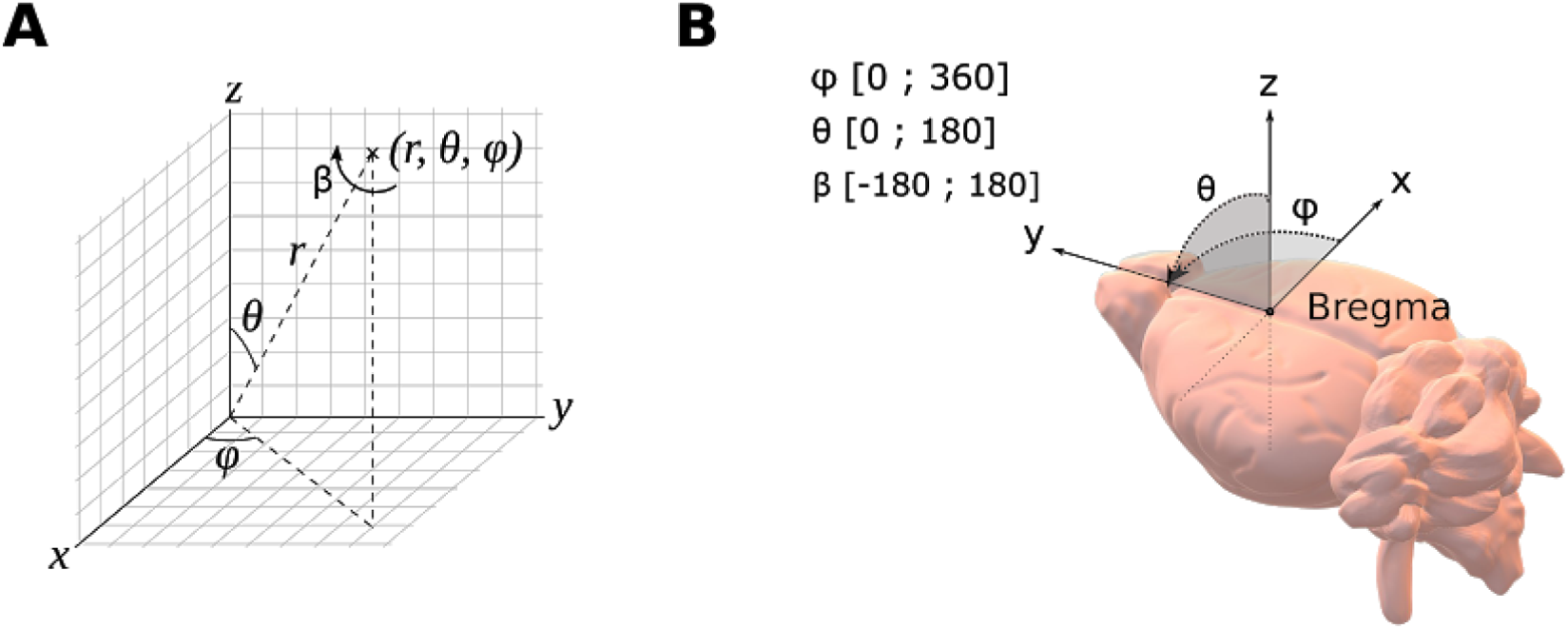

***Coordinate System: A**) Standard polar angles are used, alongside the standard [x,y,z] base. In total, an electrode trajectory registers 7 entries (3 angles, 3 coordinates, depth) to define a probe insertion. **B**) The coordinate system is aligned to the ML (x, Right: positive), AP (y, A: positive), DV (z, D: positive) stereotaxic axis and zeroed at Bregma. The bounds for each angle are indicated.*

