## Appendix 5: Alyx User Guide for "Data architecture for a large-scale neuroscience collaboration"

### Appendix 5 - IBL Alyx user guide

#### Table of contents

**General notes** 3

**Logging in** 4

**Alyx Dataflow** 5

**Navigating the database** 6
- **Subject view** 6
- **Session view** 6
- **Saving manual entries** 6

**Logging subject metadata** 7
- **Adding a new subject** 7
  - Making a subject request 8
- **Project** 8
  - Project registration 8
  - Create a project on Alyx [ADMIN] 8
  - Change project 9
  - Practice and test projects 9
- **Weighings** 10
- **Habituation and Handling** 10
- **Water restrictions** 10
- **Water supplements** 11
- **Surgeries** 12

**Cage information** 14
- **Through the Subject admin interface** 14
- **Through the Housing admin interface** 15

**Stock maintenance** 17
- **New breeding pair** 17
- **New litter** 17
- **Make new subjects for the litter** 17
- **Mark mice to be genotyped** 17
- **Give ear-marks and record that a genotype test has been done** 18
- **Record the results of the genotype test** 18
- **Mark mice to be culled** 18

**Record that mice were culled** **18**
Culling reasons - Explanatory examples 19
**Fulfilling a subject request** **20**
**Adding images to notes** **21**
**Permission system** **22**
**Notifications** **22**
**Responsible user has changed** **22**
**Water to give to mouse** **22**
**Mouse is underweight** **22**
**Summary of session datasets** **24**
**Programmatic data entry, query and backup** **25**
**Data entry** **25**
**Data query** **25**
**Backup** **25**
**Report a problem** **26**
**References** **26**

#### General notes

In this document, we will refer to a fictional lab named **yourlab.** Most links are thus fictional, and will not work if entered in a browser.

Most of the paths are given in this document as the top-right navigation bar path, for example:

- *Home › Misc › Labs*

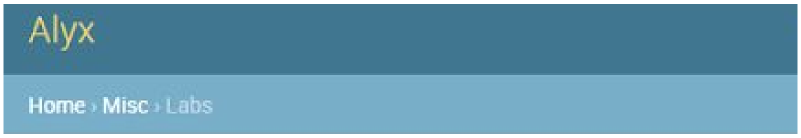


- ***This is not equivalent to the display on the Home page**, for example Labs is found under IT admin:*

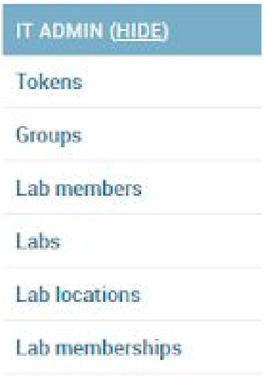


- By default, the home page should have reflected the page hierarchy, however some have expressed a preference for a more custom organization, with the most frequently used items at the top. Discussion on the Home page format is ongoing, see https://github.com/cortex-lab/alyx/issues/486

#### Logging in

- Go onto the Alyx root API
  - For the IBL: https://alyx.internationalbrainlab.org/
  - For the fictional lab: https://alyx.**yourlab**.org/ (NB: this link will not work)
- Click to join the admin interface (you should notice the link changing to .../admin) and log in.

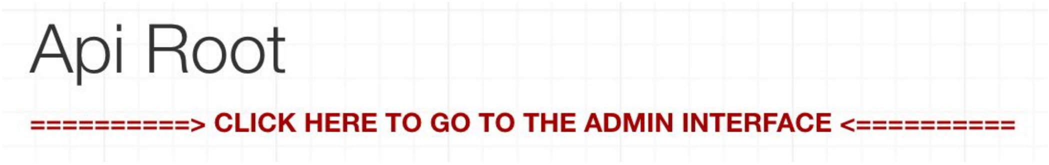


- The admin interface allows you to view the data through a web browser.
- Note that you do not require admin rights to access the admin interface, but only a regular username and password.
- Your username and password are provided by administrators (ask Olivier Winter).

Note: If you want to **test** using the website, you can open instead the development version (onto which data is deleted everyday): https://**dev.**alyx.yourlab.org/

#### Alyx Dataflow

After acquisition:

- On the rig computer:
  - Session is **created** on Alyx (if this fails, this will be done later at registration)
  - Data is **transferred** to the local server
- On the local server:
  - Data is **extracted**: from the raw session data, standardized ALF files are created
  - Data is **registered:** datasets are created in Alyx attached to the session
- Alyx then initiates the transfer from the local server Globus endpoint to the FlatIron Globus endpoint automatically.

Users can also access Alyx directly and input metadata.

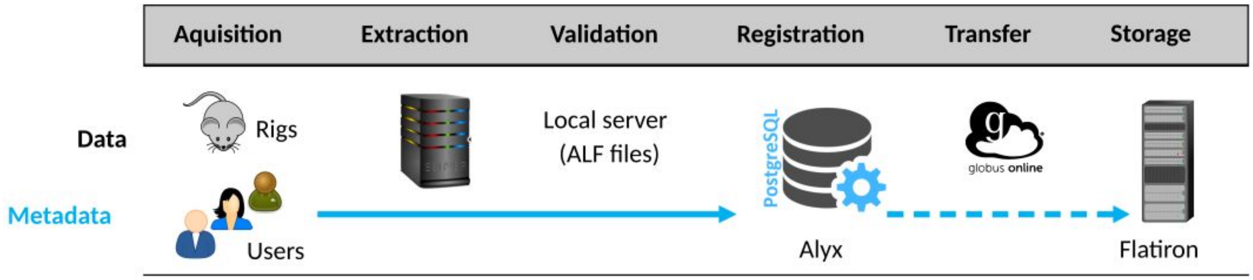

### Navigating the database

#### *Subject view*

- Click "Subjects" to see your current mice.
- Use the filters on the right to view mice that are dead, or that are in stock. You should be able to see the genotype and various other details of your mice here.
- Click on a mouse name to see all the details known to the database about that mouse. This includes the weight curves.
- From the home page, click on "Adverse Effects" to see all mice that have adverse effects recorded.

#### *Session view*

- Click on “Sessions”. The list interface shows a count of datasets with colour-code:
  - Nothing means datasets have not been registered
  - Gray means datasets have been registered but not yet transferred to the Flat Iron
  - Green means datasets are on the FlatIron server

| | | | | | | |
|---|---|---|---|---|---|---|
| IBL_13 | April 8, 2019, 10:42 a.m. | 1 | churchlandlab | - | _iblrig_tasks_biasedChoiceWorld4.0.1 | valeria |
| IBL-T3 | April 8, 2019, 9:36 a.m. | 1 | angelakilab | 27 | _iblrig_tasks_trainingChoiceWorld4.0.1 | jeanpaul |
| IBL-T1 | April 8, 2019, 9:32 a.m. | 1 | angelakilab | 25 | _iblrig_tasks_biasedChoiceWorld4.0.1 | jeanpaul |
| ibl_witten_06 | April 5, 2019, 6:49 p.m. | 1 | wittenlab | 25 | _iblrig_tasks_biasedChoiceWorld4.0.1 | alejandro |
| ibl_witten_05 | April 5, 2019, 6:46 p.m. | 1 | wittenlab | 25 | _iblrig_tasks_biasedChoiceWorld4.0.1 | alejandro |
| ibl_witten_04 | April 5, 2019, 6:45 p.m. | 1 | wittenlab | 25 | _iblrig_tasks_biasedChoiceWorld4.0.1 | alejandro |
| dop_3 | April 5, 2019, 5:38 p.m. | 1 | wittenlab | 27 | _iblrig_tasks_trainingChoiceWorld4.0.1 | alejandro |
| dop_2 | April 5, 2019, 5:23 p.m. | 1 | wittenlab | 27 | _iblrig_tasks_trainingChoiceWorld4.0.1 | alejandro |
| dop_1 | April 5, 2019, 5:19 p.m. | 1 | wittenlab | 27 | _iblrig_tasks_trainingChoiceWorld4.0.1 | alejandro |

1 2 3 4 ... 87 88 4360 sessions

#### *Saving manual entries*

Make sure to click "save" (or press enter) every time you make a change that you want to be saved.

In many cases you can use the "save and continue editing" button to save but stay on the page to enter more information.

#### Logging subject metadata

**Important note**:

- Make sure to click "**save**" (or press enter) every time you make a change that you want to be saved.
- In many cases you can use the "**save and continue editing**" button to save but stay on the page to enter more information.

##### *Adding a new subject*

**Mandatory Immediately:**

- **Sex** (Subject tab)
- **Birth date** (Subject tab)
- **Lab** (Subject tab)
- **Project** (see section below for details)

**Mandatory:**

- **Ear mark** (Subject tab)

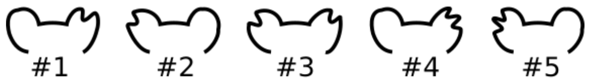


    - IBL nomenclature: R L RL RR LL

- **Cage** (Profile tab)
    - The cage name/number can be arbitrarily given if not enforced within the institution.
- **Strain** (Profile tab) (mouse's background)
- **Line** (Profile tab) (e.g. transgenic line)
    - For a Wild Type mouse, put same info for Strain and Line (see image below)

- 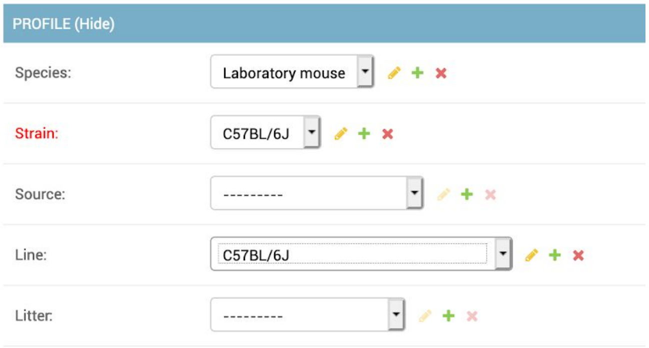


- **Litter**
  - For each new litter ordered from Jax, first go to main/Litters and create a new litter (e.g. CSHL_Jax_001) leaving the breeding pair blank. Then add this for each animal when entering it into Alyx.

##### *Making a subject request*

Making subject requests when you need mice helps the lab manager plan breeding pairs and helps distribute mice fairly.

To make a subject request, from the home page click "add" to the right of "subject requests" and specify the fields as suggested. Use the notes field to specify any requirements not otherwise listed, such as gender or age of the mice you need.

When the manager has fulfilled the request, you will get an email (assuming you have set your email address) and the mice will show up in your Subjects view.

#### ***Project***

##### *Project registration*

Prior to adding your mice onto the database, you should have registered and created a project.

In the IBL, a project is registered on GitHub (i.e. described and available to view for the community) via issues on a dedicated repository. However, any platform could be used to do so. Once a project is registered on a platform, it has to be created on Alyx.

Note: **Only administrators can create projects. Contact an admin if you need to add a project to the database.** Steps to add a project are given below.

##### *Create a project on Alyx [ADMIN]*

- To create a project, Go to Other>Projects, click on +
- The IBL uses the following naming convention :
    - **`ibl_keywords`** for **general collaborative projects**
        - e.g. `ibl_neuropixel_brainwide_01`
        - `ibl_` is at the beginning.
    - **`labname_keywords`** for **personal projects**
        - e.g. `witten_dop`
    - The project name is **unique** (enforced by the database).
- Make sure to add the GitHub link (or any other platform used) in the project description.
- Select users as appropriate (project leads and collaborators)

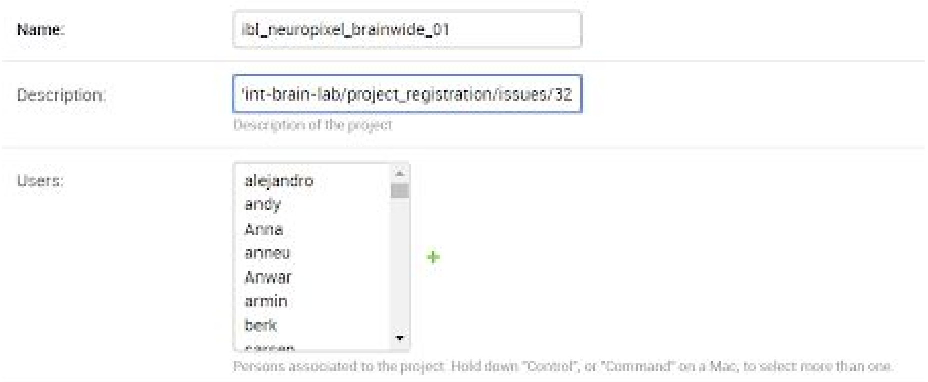


- If following the IBL model, write in a comment on the corresponding GitHub issue:
    - “ Now registered as *<project code name>* on Alyx “

#### *Change project*

A mouse could be attributed to a (or several) different project(s). To do so :

- Go onto the *Subject* page, and select the new appropriate project(s)

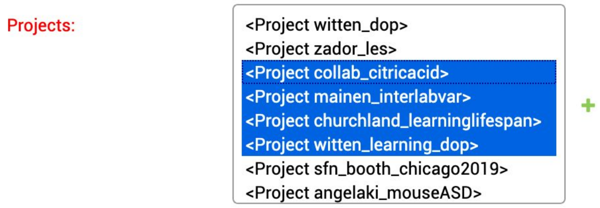


- Those changes will be propagated to the future sessions acquired (not retrospectively).
    - If you need to retrospectively change the project for a session, you can do so in the *Session* page.

#### *Practice and test projects*

You may need to do some procedural practice, or some test. In which case, you may not want to use the main project code (in order to facilitate down-the-line analysis).

It is suggested to use the project code:

- **Practice** : When the main main project guidelines are followed, with some minor divergences (e.g. if the animal is older than what was previously intended). The data collected is intended to be on the main pipeline, but the Practice project tag serves as a flag.
    - Add **Practice** to any other current project code on the Subject page.

- **Test** : When out-of-the-pipeline tests procedures are performed. The data collected is not intended to be part of the main analysis pipeline.
    - Add **Test** and <u>remove</u> any other current project code on the Subject page.

Note: This is not implemented per session, but per subject. Ongoing discussion can be found at https://github.com/cortex-lab/alyx/issues/606

##### ***Weighings***

- From the home page, click the "add" button to the right of "weighings" and fill in the details.
- Weighings performed with pyBpod software should be automatically logged so no need to enter manually.

##### ***Habituation and Handling***

- In subject page, go to 'Other action' tab and select `handling_habituation` under 'procedure'.
- Write session narrative.

*Note*: If you cannot find a procedure, you will need to create it: *Home › Action admin › Procedure types › Add procedure type* https://alyx.yourlab.org/admin/actions/proceduretype/add/ . This procedure will be visible to all mice in yourlab automatically, so the procedure needs only to be created once.

##### ***Water restrictions***

- Use the "add" button next to water restrictions to put a mouse on water restriction.
- The reference weighing will be the last one before the start time. This should be logged in as any other weighing from the "Weighings" page.
- When beginning a water restriction remember to weigh the implant and the mouse separately. The fields will ask you for both to take into account a baseline. **Note:** This

reference weight should be the same as the weighing you added on the “Weighings” page before creating a water restriction.

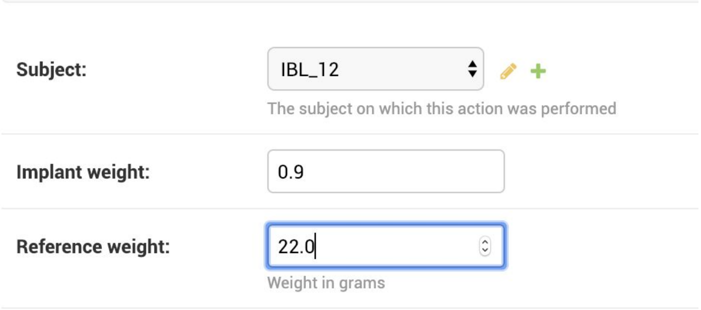


- To take a mouse off water restriction, give an end date to the water restriction that already exists:
    - go to 'Water Restrictions'
    - click on the 'Start Date' for the subject of interest
    - fill the end time (date and time)
    - Save
- Then, to put the mouse BACK on water restriction, create another new water restriction.
- See section ***Notification - Mouse is underweight*** to set the weight threshold for mice on water restriction.

##### *Water supplements*

- From the home page, click the “Water administrations” button and click on the “add water administration” button to create a new water administration.
- Select a subject from the drop-down menu and fill in the date and time details. **Note** that you are able to create administrations for the future by simply indicating a future date and time.
- In the “Water administered:” field, place the number of milliliters you administered if your water administration was a controlled amount. For instance, the amount below equals to 1 mL.
- Indicate the type of water you are administering to the mouse from the drop down menu labeled “Water type:”

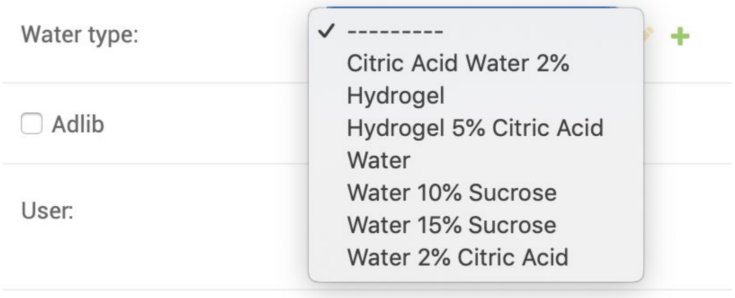


- If you are giving the mouse ad lib water of any kind, click on the Adlib button to indicate this. **Note:** Alyx does not indicate that the mice have ad lib water for future days. For instance, if you are giving ad lib water, hydrogel, etc. on a Friday until Sunday, if you create this water administration on Friday it will only be recorded for that day. You are then able to create future ones for the rest of the weekend or days needed.

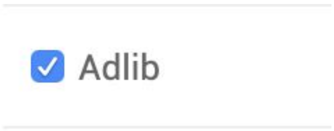


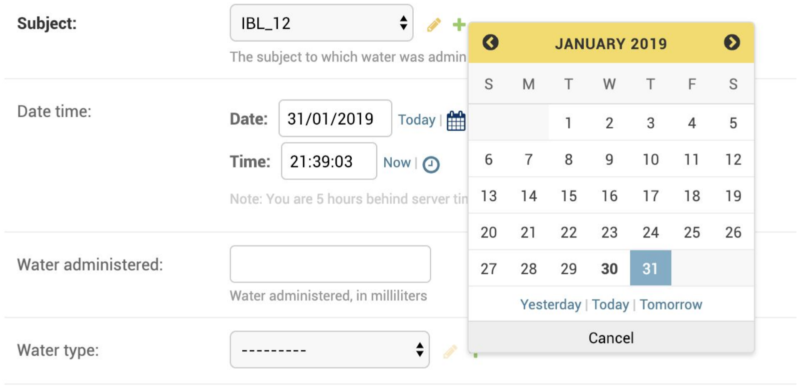


#### *Surgeries*

- From the home page, click the "add" button to the right of "Surgeries".
- Choose your mouse's name from the dropdown and fill in any relevant details (example below).

- Note: if the mouse you're looking for does not appear in the dropdown, it is because you are not the responsible user for that mouse.
  - If you intend to be, then first "get the mouse from stock" - see below.
  - If you are entering a surgery for someone else's mouse, go to the mouse's subject page and find the surgeries section near the bottom. Add the surgery there, then click "save and continue editing" to get an orange "edit" icon for that

surgery - click it to bring up the full page for that surgery to enter any further details.

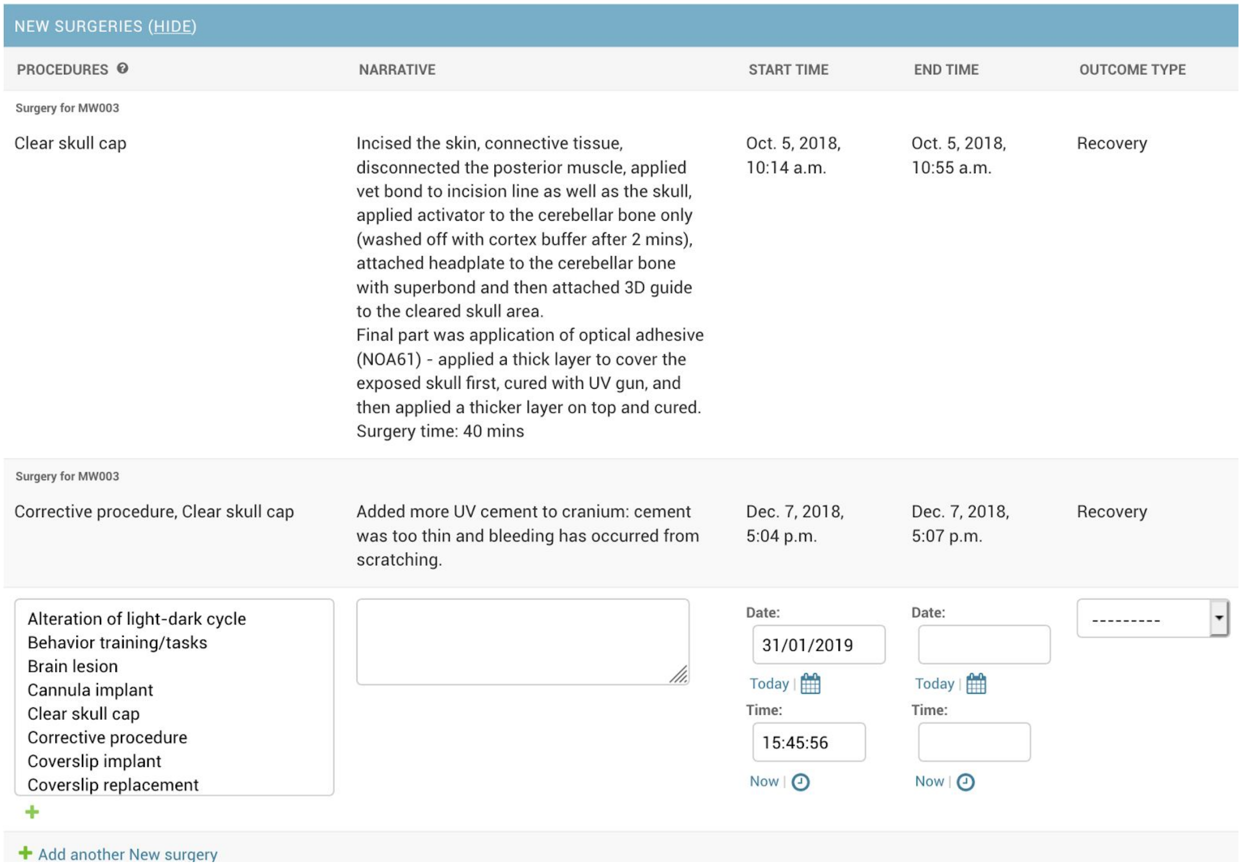

#### Cage information

- When creating a *Housing* object, the default values will be populated from your user current lab. You can set your default lab housing conditions through the Lab admin form located at *Home › Misc › Labs* https://alyx.yourlab.org/admin/misc/lab/
- Since we need to keep history of housing conditions, you have to assign a mouse to a cage explicitly through a *HousingSubject*. You can think of it as a home lease.
- Since we have 3 models: *Housing, Subject* and *HousingSubject* there are several ways to access/update cage information below.

##### *Through the Subject admin interface*

- Using the top right navigation bar, go to the subject page *Home › Subject › Subjects*
- Go to the bottom of the page under the *Housing Subjects* page
- Create and/or modifiy a *HousingSubject*, which assigns an Housing to a Subject. It is important to set the start date properly so that the history is properly recorded.
    - Example of creation (click on + Add another House subject):

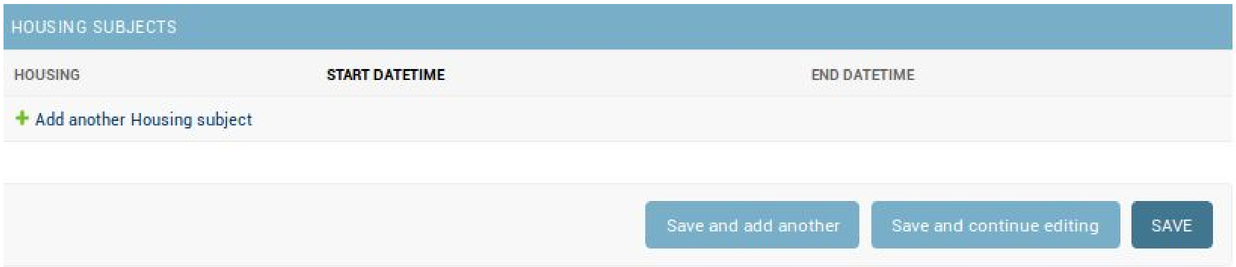


- If a new cage is used, click on + and add cage information (then Save):
    - **Cage name** is a unique ID (as given at your institution), that can be shared across co-housed mice

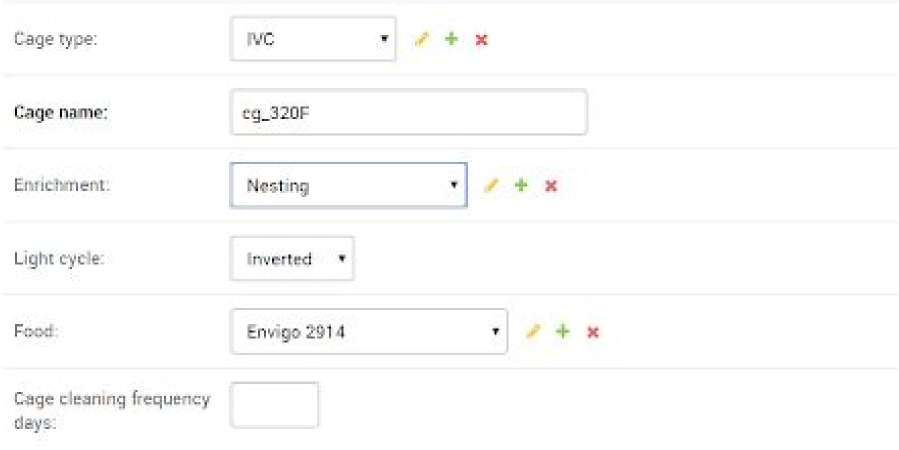


- Example of assigning an existing cage:

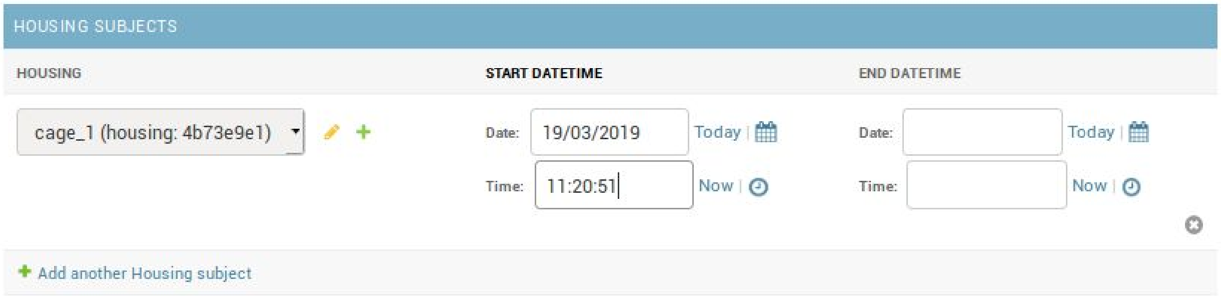


- You can also create a *Housing* object from here.

#### ***Through the Housing admin interface***

- Using the top right navigation bar, go to the subject page *Home › Misc › Subjects > Housing*
- Fill up the fields using listboxes. If items are missing, you can create them.
- You can assign mice to this new *Housing* through the *HousingSubjects* widget at the bottom of the page

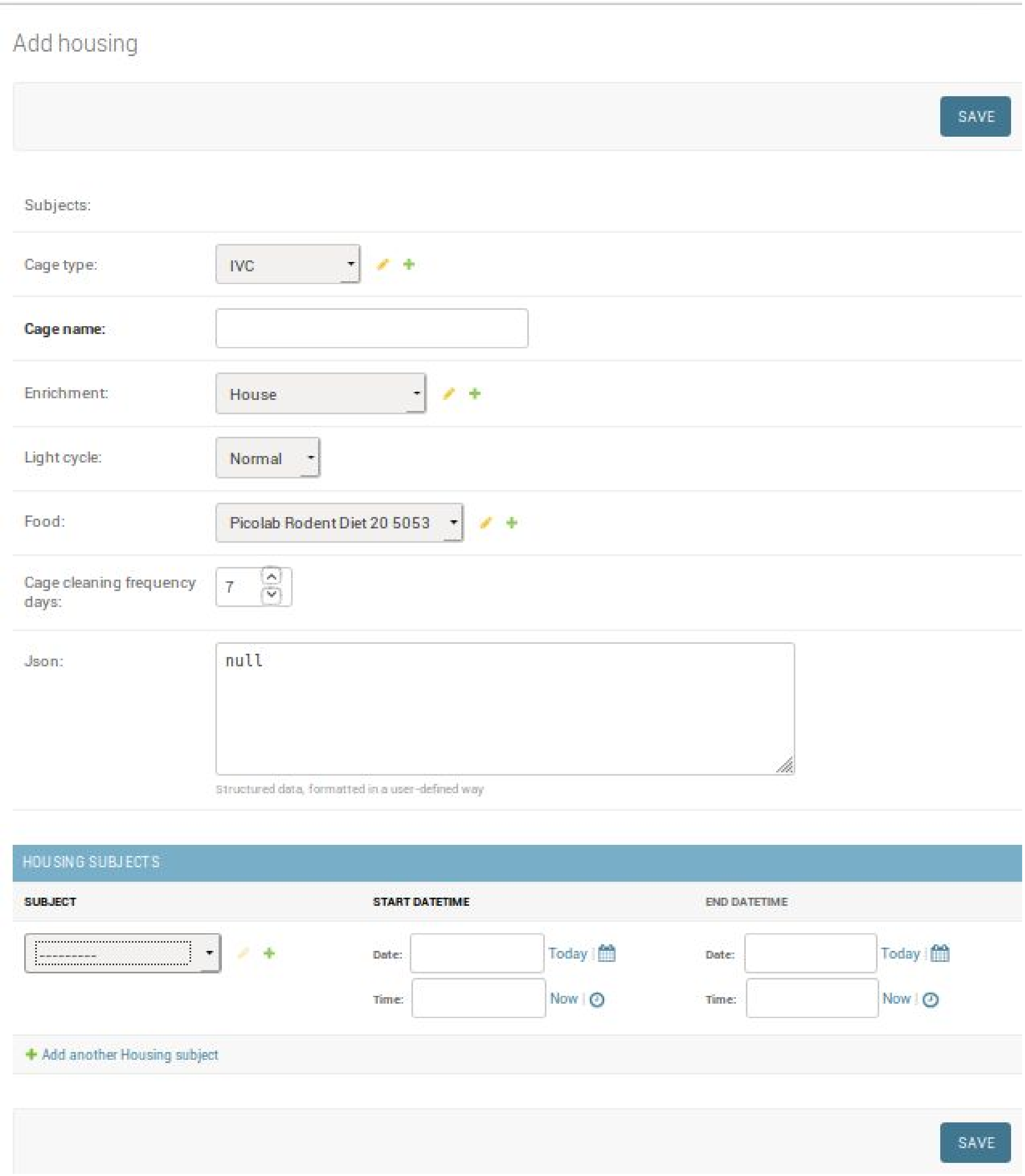
Add housing
SAVE
Subjects:
Cage type:
IVC
Cage name:
Enrichment:
House
Light cycle:
Normal
Food:
Picolab Rodent Diet 20 5053
Cage cleaning frequency days:
7
Json:
null
Structured data, formatted in a user-defined way
HOUSING SUBJECTS
SUBJECT
START DATETIME
END DATETIME
---------
Date:
Today
Time:
Now
Date:
Today
Time:
Now
Add another Housing subject
SAVE

#### Stock maintenance

##### *New breeding pair*

- From the home page, click "add" to the right of "Breeding Pairs". Select the line of mouse with the dropdown, then click "save and continue editing" at the bottom. The name of the breeding pair will be automatically generated.
- Add a start date, and choose the Father and Mother(s). You need to know the mice you'd like to use by number. These dropdowns only show living male or female mice from this particular line, so if you don't see the mouse you want, you may need to go to that mouse's subject page and change its line to match the line for which you would like it to be the breeding parent.
- Click Save when finished.

##### *New litter*

- From the home page, click "add" to the right of "Litters".
- Select the line of the mice with the dropdown then click "save and continue editing" at the bottom. The name of the litter will be automatically generated.
- Choose the birth date and the breeding pair, and click save.

##### *Make new subjects for the litter*

- When the pups are weaned, you can create individual entries for the mice in the litter with their genders and wean date. To do this, go to the editing page for the litter, which you can get to either by :
    - clicking "Litters" from the home page, sorting by line with the filter to the right, and finding the litter in question (if you know the litter number);
    - OR by first going to the Breeding Pair by clicking "Breeding Pairs", sorting by line, choosing the breeding pair in question, then clicking the orange "change" button in the list of that BP's litters near the bottom of the page.
- Once you are at the litter edit page, scroll down to the list of subjects (so far empty), and select the gender for the first pup.
- Click "Save and continue editing" to see this new pup get an automatically generated name.
- Repeat for each additional pup.
- Click "Today" for all the Wean Date fields and click Save.

##### *Mark mice to be genotyped*

- Return to the litter page, as above, and select the To Be Genotyped checkbox for all the mice to be genotyped.
- Click Save.

##### *Give ear-marks and record that a genotype test has been done*

- Return to the litter page as above. Click "Today" for all the genotype date fields, and enter the ear marks in the field (tip: to go from one mouse's ear mark field straight to the next, use the down arrow key).
- Click Save.

##### *Record the results of the genotype test*

- Return to the litter page as above and find the columns for Sequence and Result.
- For each sequence that was tested, choose whether the result for that mouse was Absent or Present.
- Click Save.
- When you view these mice in the Subjects view, their zygosities for the relevant alleles should have been automatically calculated.

##### *Mark mice to be culled*

- From the home page, click "Cull subjects". Filter the view as necessary and select "To be culled" checkboxes.
- Click save.

##### *Record that mice were culled*

- If you cull your mouse, go to the subject page for that mouse and record its Death date (Reduced date is only relevant for the UCL database - ignore the field).

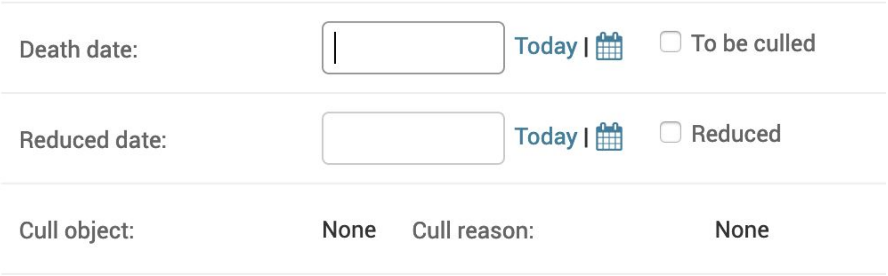


    - Note: you need to be the responsible user or an admin in order to cull an animal.
    - You can also access the list of subjects to cull via this page: *Home › Subject admin › Cull_subjects* https://alyx.yourlab.org/admin/subjects/cull_subject/ Filter the view as necessary and click "Today" for the death date of any mice you culled, or that were culled.

- Once a date has been entered, a Cull object appears.

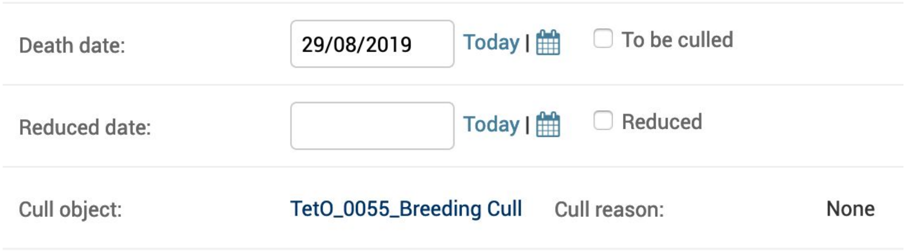


- ○ Click on the Cull object.
  See section ***Culling reasons - Explanatory examples*** if you are unsure on what to select.

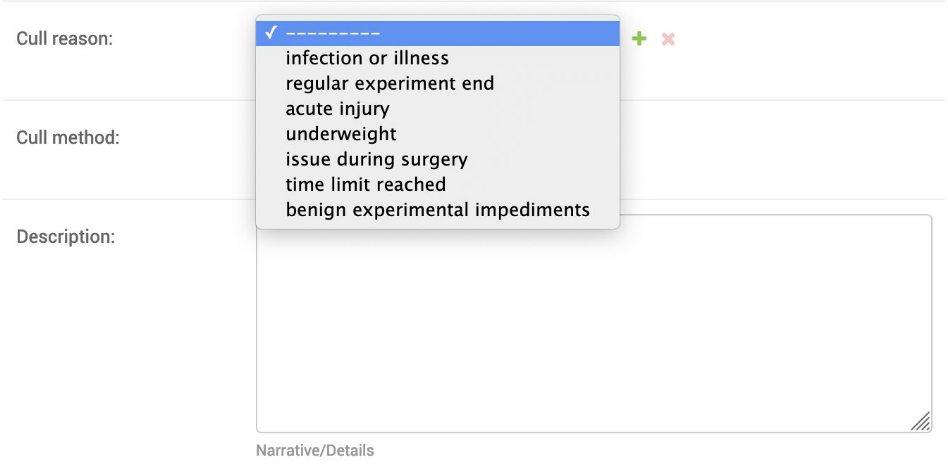


- ○ Enter
  - ■ Cull reason
  - ■ Cull method
  - ■ Description (describe reason for culling if not anticipated)

- Then fill in the fields under the "outcomes" section: Cull method, Adverse effects, and Actual Severity.
- You can view all mice and associated cull reason through this web page on Alyx: *Home > Action admin > Culls* https://alyx.yourlab.org/admin/actions/cull/

- If you want to add pictures related to your perfusion, create a new 'Surgery' (Non-recovery, 'Perfusion' as type of surgery) and add photos there.

*Culling reasons - Explanatory examples*

- **Infection or illness:** When the animal cannot be treated and recovered.
- **Regular experiment end:** When everything went according to plan.
- **Acute injury:** e.g. loss of implant, animal found dead in cage.

- **Underweight:** When severity limit is reached and animal has to be culled.
- **Issue during surgery:** e.g. if consequent bleeding occurs during a craniotomy and the animal has to be sacrificed.
- **Time limit reached:** e.g. if the protocol only allows to experiment on subject for a certain duration.
- **Benign experiment impediment:** e.g. if the animal develops a cataract.

***Fulfilling a subject request***

- From the home page, click Subjects and filter the view as necessary (e.g. mice that are alive, in a certain line, in Stock, have all zygosities positive).
- Change the responsible user of the mice to match the user who requested them.
- Click Save.

#### Adding images to notes

- Select a subject from the “Subjects” page on the home screen
- Select the Notes section
- Input the responsible user, date and time, and fill out the description.
- Next to the ‘TEXT’ box select the ‘Choose File’ button

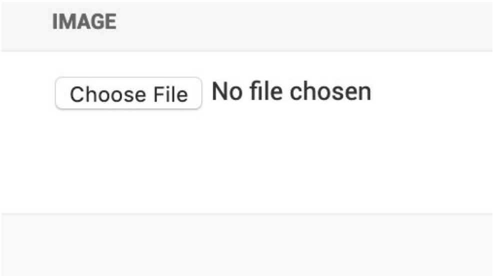


- Choose your file and upload, then click ‘Save’,‘Save and continue editing’, or ‘Save and add another’.

#### Permission system

- Superusers can modify all subjects.
- Other users can only modify their own subjects and the data associated to them (surgeries, water restrictions…).
- Delegation: User A can add Users B and C to his "allowed_users" field in their profile page, so that Users B and C can modify User A's subjects.

#### Notifications

Please make sure you input your email address appropriately in: *Home > Misc > Lab members* https://alyx.yourlab.org/admin/misc/labmember/

Only one email address can be entered. This address will be used to send emails containing notifications. Types of notifications outlined below.

You can control the notifications you receive here: *Home › Action admin › Notification rules › Add notification rule* https://alyx.yourlab.org/admin/actions/notificationrule/

You can see all notifications pending in and already sent: *Home › Action admin › Notifications* (select 'All' and the type of notification) https://alyx.yourlab.org/admin/actions/notification/

##### *Responsible user has changed*

An email is sent to the user if:

- The responsible user for an animal changes

##### *Water to give to mouse*

An email is sent to the user if:

- A water administration has not been logged on Alyx within the last 23 hours
- If extra water needs to be given to the animal on the day.

This cron job is run hourly.
https://github.com/cortex-lab/alyx/blob/master/alyx/actions/notifications.py#L33

##### *Mouse is underweight*

An email is sent to the user if:

- Sends a 'Warning' if the weight drops below +2% of the set threshold
- Sends a 'WARNING' of the weight drops below the set threshold

https://github.com/cortex-lab/alyx/blob/master/alyx/actions/notifications.py#L18

The threshold can be set in, however only by admin users : *Home › Misc › Labs*
https://alyx.yourlab.org/admin/misc/lab/
Select your laboratory, and change Reference/Zscore weight percentage (pct) according to your protocol:

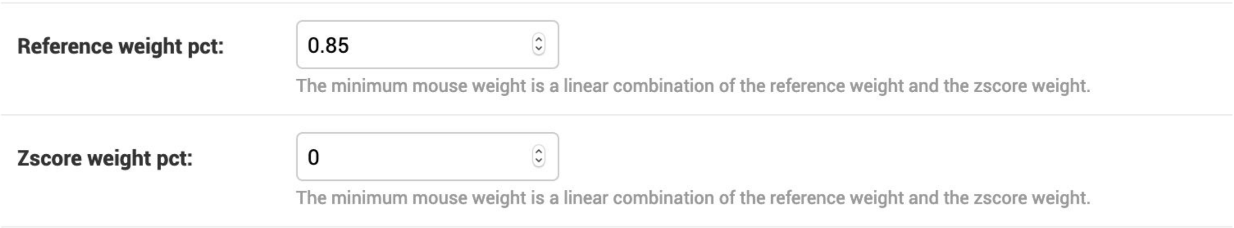

#### Summary of session datasets

To check your data is online (i.e. uploaded to FlatIron), go to Home > Action admin > Sessions. See column #DATASETS.

- A **dash** ( - ) means no data has been registered to the session
- A **gray number** means data has been registered, but not uploaded to FlatIron or partially uploaded
- A **green number** means everything is online

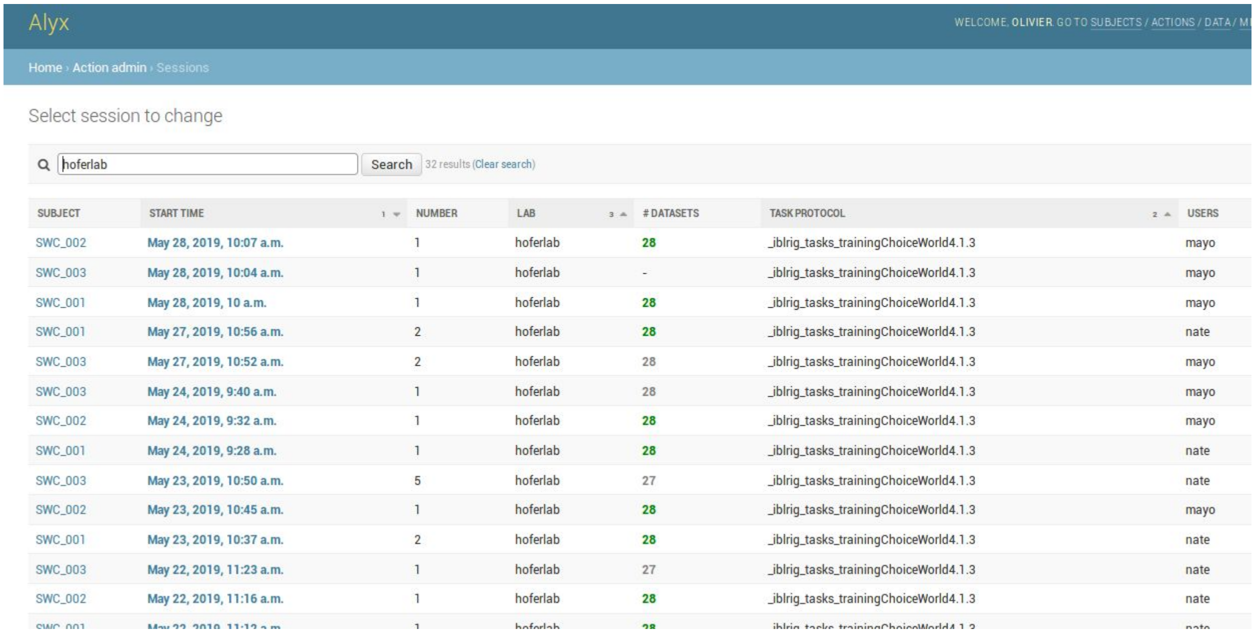


| SUBJECT | START TIME | NUMBER | LAB | # DATASETS | TASK PROTOCOL | USERS |
|---|---|---|---|---|---|---|
| SWC_002 | May 28, 2019, 10:07 a.m. | 1 | hoferlab | 28 | _iblrig_tasks_trainingChoiceWorld4.1.3 | mayo |
| SWC_003 | May 28, 2019, 10:04 a.m. | 1 | hoferlab | - | _iblrig_tasks_trainingChoiceWorld4.1.3 | mayo |
| SWC_001 | May 28, 2019, 10 a.m. | 1 | hoferlab | 28 | _iblrig_tasks_trainingChoiceWorld4.1.3 | mayo |
| SWC_001 | May 27, 2019, 10:56 a.m. | 2 | hoferlab | 28 | _iblrig_tasks_trainingChoiceWorld4.1.3 | nate |
| SWC_003 | May 27, 2019, 10:52 a.m. | 2 | hoferlab | 28 | _iblrig_tasks_trainingChoiceWorld4.1.3 | mayo |
| SWC_003 | May 24, 2019, 9:40 a.m. | 1 | hoferlab | 28 | _iblrig_tasks_trainingChoiceWorld4.1.3 | mayo |
| SWC_002 | May 24, 2019, 9:32 a.m. | 1 | hoferlab | 28 | _iblrig_tasks_trainingChoiceWorld4.1.3 | mayo |
| SWC_001 | May 24, 2019, 9:28 a.m. | 1 | hoferlab | 28 | _iblrig_tasks_trainingChoiceWorld4.1.3 | nate |
| SWC_003 | May 23, 2019, 10:50 a.m. | 5 | hoferlab | 27 | _iblrig_tasks_trainingChoiceWorld4.1.3 | nate |
| SWC_002 | May 23, 2019, 10:45 a.m. | 1 | hoferlab | 28 | _iblrig_tasks_trainingChoiceWorld4.1.3 | mayo |
| SWC_001 | May 23, 2019, 10:37 a.m. | 2 | hoferlab | 28 | _iblrig_tasks_trainingChoiceWorld4.1.3 | nate |
| SWC_003 | May 22, 2019, 11:23 a.m. | 1 | hoferlab | 27 | _iblrig_tasks_trainingChoiceWorld4.1.3 | nate |
| SWC_002 | May 22, 2019, 11:16 a.m. | 1 | hoferlab | 28 | _iblrig_tasks_trainingChoiceWorld4.1.3 | nate |
| [illegible] | [illegible] | 1 | [illegible] | [illegible] | [illegible] | [illegible] |

You can also get the information on a datasets by typing the UUID after the link [https://alyx.yourlab.org/datasets/**[UUID]**](https://alyx.yourlab.org/datasets/[UUID])

For example: [https://alyx.yourlab.org/datasets/a9d025e9-9ac9-49d0-a9c8-bf156873278f](https://alyx.yourlab.org/datasets/a9d025e9-9ac9-49d0-a9c8-bf156873278f)

#### Programmatic data entry, query and backup

##### *Data entry*

A "REST API" is available at alyx.yourlab.net. You can use this to read data in an easy and conveniently-formatted way, and to post certain kinds of data like weighings and water administrations.

See the alyx-matlab github repository's "Examples.m" for more about how to do this via Matlab, and https://github.com/int-brain-lab/ibllib/tree/master/examples/oneibl for an example in Python using ONE.

##### *Data query*

Queries can also be performed with the REST API, using the Alyx.getData method in the github repository.

SQL queries can also be performed directly using the JDBC connection to the Alyx's postgresql database. See exampleSQLquery.m.

##### *Backup*

One can set up a cron job to make automated daily SQL and JSON backups.

#### Report a problem

Go to GitHub https://github.com/cortex-lab/alyx/issues to report a problem or ask a question.
